# *Arhopalus rusticus* (Coleoptera: Cerambycidae): a new vector for pine wood nematode, *Bursaphelenchus xylophilus* (Nematoda: Aphelenchoididae)

**DOI:** 10.1101/2020.06.09.142588

**Authors:** Yang Wang, Yunfei Mao, Fengmao Chen, Lichao Wang, Juan song, Lifeng Zhou

## Abstract

In the study on the transmission of pine wood nematodes by *Arhopalus rusticus*, we research the ability and characteristics of *Arhopalus rusticus* to transmit pine wood nematodes. A total of 288 *Arhopalus rusticus* (female 142, male 146) were collected and 20 (female 8, male 12) of them were able to carry pine wood nematodes. Results shown that 12 beetles can transmit pine wood nematodes. *Arhopalus rusticus* does not eat the pine bark, but eat the pine needles. Pine wood nematodes are transmitted through the feeding bite mark on the pine needles. In addition experiments further confirmed that pine wood nematodes could transmit through pine needles.

## 1 Introduction

Pine wilt disease (PWD) caused by the pine wood nematode (PWN), *Bursaphelenchus xylophilus*, is the most destructive disease that affects pine species and it causes significant economic losses [1]. Pine wilt disease (PWD) is regarded as “pine cancer” because the infected pines will die rapidly and no effective measures are available for its treatment. Pine wood nematode(PWN) is considered to be native to North America, where it usually only damages exotic pine trees, but it has also spread to Asian and European countries such as Japan, China, South Korea, Portugal, and Spain[2, 3]. The spread of PWD is rapid and it is extremely difficult to prevent and control. Therefore, PWD has been listed as a quarantine object in many countries.

Studies have shown at least 45 insect species can carry PWN, which they belong to Cerambycidae, Buprestidae, Curculionidae, Scolytidae, and Termitidae. However, not all insects that carry PWN can transmit it and only insects with life histories that match that of PWN can become vectors. The activities of vector insects, especially supplemental nutrition and spawning, are the basic modes responsible for the natural transmission of PWN[4]. Thus, among the 45 insects that carry the PWN, only 13 can be used as vectors. These 13 insect species all belong to the family Cerambycidae (*Monochamus* spp.). The regional distributions of these 13 beetle species comprise *Monochamus alternatus, Monochamus saltuarius, Monochamus grandis*, and *Monochamus sutor* distributed in Asia, *Monochamus galloprobincialis* and *M. sutor* distributed in Europe, and eight beetle species distributed in North America [5-8].

### Whether the other carrying-nematodes beetles could spread nematode?

*Arhopalus rusticus* (Coleoptera: Cerambycidae), Genus: *Arhopalus*. It is an important wood-boring pests. Known host records show that *Arhopalus* is mainly associated with coniferous plants, particularly Pinaceae and Taxodiacea and debilitated wood of variety of conifers[9]. *Arhopalus* is a Northern Hemisphere cerambycid genus with about 25 species and subspecies and now occurs in all major biogeographic regions of the world through the spread of commerce [9-11]. *Arhopalus rusticus* is the most powerful wood-boring pest carrying *Bursaphelenchus mucronatus* besides the *Monochamus alternatus*, but it is not clear whether they can carry and spread pine wood nematodes [12]. Linit et al. listed *A. rusticus* carrying *Bursaphelenchus xylophilus* larvae in Japan and North America [6, 13]. M Jurc et al. believes that brown beetle is a potential vectors of *Bursaphelenchus xylophilus*[14]. Kulinich et al. thought in Russia, *A. rusticus* was observed to act as a carrier of *Bursaphelenchus spp*[15]. Y Mamiy et al. detected pine wood nematode from *Arhopalus rusticus* and believed that *Arhopalus rusticus* may have the possibility of spreading *Bursaphelenchus xylophilus*. [16, 17].

Although the above studies suggest that *Arhopalus rusticus* can carry PWN and is a potential medium, there is no study to indicate that it can transmit PWN. In this study, we study whether *Arhopalus rusticus* could transmit PWN.

## 2 Materials and methods

### Part 1 Population survey of *Arhopalus rusticus* in forests and the situation of they carry PWN

Between May and September during 2017, beetles were collected from Qingdao in Shandong province and Dalian in Liaoning province, China. APF-I type attractants were used to trap beetles in pine wilt disease (PWD) epidemic areas. The numbers and species of the beetles were recorded Every day, and PWN in the beetles were detected by the Baermann funnel method[18]. The species of nematodes was identified with a microscope (Zeiss Axio Lab.A1).

### Part 2 Study on the transmission of PWN by *Arhopalus rusticus*

#### Collection of Arhopalus rusticus

Collected dead trees of *Pinus massoniana* from December 2017 to March 2018 in Qingdao, Shandong province, China. The trees which were infested by PWNs and *Arhopalus rusticus* larvae, were cut into logs (1.5 – 2m) and maintained in a large field cage (2.0 m × 2.0 m × 2.0 m) with 0.8 mm mesh wires netting. *Arhopalus rusticus* were collected daily during adult emerge (from May 10th to July 11th)from the logs. In this study, 288 *Arhopalus rusticus* were collected for analysis the transmission characteristics of the PWN through adult feeding.

#### Rearing method of Arhopalus rusticus

Newly emerged beetles were collected daily and feeding individually in insect bottle (h = 20cm, r = 4cm) with one-year-old fresh and healthy twigs(diameter:0.5-1cm, length:10-15cm) of *Pinus massoniana*. In each insect bottle only one *Arhopalus rusticus* (Male and female beetles are fedng separately after mating) and one twig (the incision was sealed with wax). The insect bottles were placed in the laboratory (25°C). Twigs were changed every three days.

#### Extracted and identification of PWN

Observe the beetles’ feeding and spawning and taking pictures(Zeiss SteREO Discovery.V20), the photographs of *Arhopalus rusticus* and Mouthpart are shown in the Appendix. Then cut the changed twigs into pieces. The nematodes in the twigs were extracted by the Baermann funnel method[18]. Twenty-four hours after exteact, suspension (10ml) at the bottom of the funnel was collected. The number and species of nematodes in the suspension (10ml) was identified with a microscope (Zeiss Axio Lab.A1).

#### PWNs transmission experiment of Pine needles

Insert the fresh and healthy pine needles into the vial bottles and add the appropriate amount of sterile water to the bottles, cut the top of the needles and place the tampon covered with PWNs (Fig 1). Place the vial bottle in a large glass cover to keep the air moist. After 12 hours, observe whether there is PWN in the liquid in the vial bottles under the microscope. And rinse the surface of the needles with water, then cut and use the Bellman funnel method to separate PWN in the needle and count.

**Figure 1.**
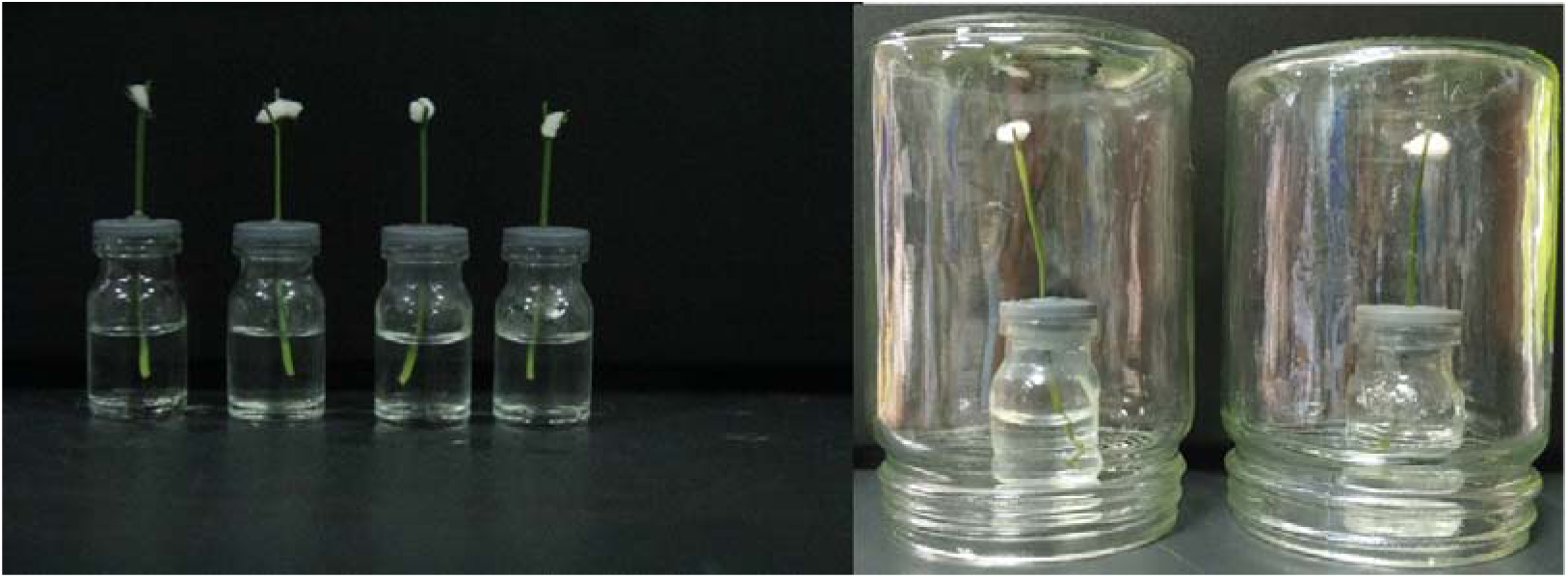
PWN transmission experiments.

Operation precautions: ➀ Fresh pine needles. ➁ Operation in time after the incision. ➂ The tampons were in full contact with the incisions. ➃ Vibrant PWNs (PWNs has just been extracted from the culture medium). ➄ Must be placed under a glass cover for moisturizing.

## 3 Results

### 3.1 Population survey of *Arhopalus rusticus* in forests and the situation of they carry PWN

The results of the investigation of the beetles in the PWD epidemic area in Qingdao in Shandong province and Dalian in Liaoning province are shown in Table 1, which demonstrates that 12 beetle species were collected in Qingdao with a total of 4223 specimens and 4 beetle species in Dalian with a total of 340. The major beetle species in PWD epidemic area were *A. rusticus* accounted for 72% of the total in Qingdao and 87% in Dalian.

**Table 1.** Beetles detected in the PWD epidemic area in Qingda and Dalian

| species of beetles | Qingdao <sup>a</sup> | Dalian <sup>b</sup> | major host plants of beetles |
| --- | --- | --- | --- |
| *Aromia bungii* | 20 |  | *Amygdalus persica*, *Armeniaca vulgaris*, *Cerasus pseudocerasus*, *Cerasus japonica*, *Salix babylonica*, *Populus L.* |
| *Anoplophora chinensis* | 1 |  | *Platanus acerifolia*, *Populus L.*, *Salix babylonica*. |
| *Acanthocinus griseus* |  | 5 | *Pinus koraiensis*, *Pinus tabuliformis*, *Pinus armandii*, *Picea asperata* |
| *Acalolepta sublusca* | 2 |  | *Buxus megistophylla*, *Euonymus alatus* |
| *Apriona rugicollis* | 1 |  | *Broussonetia papyrifera*, *Broussonetia papyrifera* |
| *Arhopalus rusticus* | 3040 | 296 | *Pinus* Linn, *Abies* Mill, *Picea*, *Larix gmelinii*,<br>*Cupressaceae*. |
| *Dorysthenes granulosus* | 1 |  | *Saccharum officinarum*, *Quercus acutissima*,<br>*Taxodiaceae*. |
| *Monochamus alternatus* | 1073 | 24 | *Pinus* Linn, *Abies* Mill, *Picea*, *Tsuga* Carr,<br>*Pseudotsuga*, *Larix*, *Cedrus*, *Morinda parvifolia*,<br>*Quercus acutissima*, *Malus pumila*, *Carthamus*<br>*tinctorius*. |
| *Monochamus saltuarius* |  | 15 | *Picea asperata* |
| *Semanotus sinoauster* | 70 |  | *Taxodiaceae*, *Cryptomeria fortunei*, *Pinus* Linn. |
| *Megopis sinica* | 12 |  | *Malus domestica*, *Crataegus pinnatifida*, *Ziziphus*<br>*jujuba*, *Diospyros kaki*, *Castanea mollissima*,<br>*Juglans regia*. |
| *Trirachys orientalis* | 1 |  | *Populus* L., *Salix babylonica*, *Sophora japonica*,<br>*Paulownia*, *Quercus acutissima*, *Quercus*<br>*acutissima*. |
| *Xylotrechus magnicollis* | 1 |  | *Ficus elastica*, *Diospyros* Linn, *Quercus*<br>*mongolica*. |
| *Xylotrechus quadripes* | 1 |  | *Coffea*, *Mangifera indica* L. *Artocarpus*<br>*heterophyllus*, *Tectona grandis*, *Catunaregam*<br>*spinosa*, *Catunaregam spinosa*, |

<sup>a</sup>: The species and quantity of beetles collected in Qingdao. <sup>b</sup>: The species and quantity of beetles collected in Dalian.

A total of 3336 *Arhopalus rusticus* were collected in this study accounted for 73.1% of the total beetles (4563). 200 *Arhopalus rusticus* were detected and 15 of them were able to carry PWN. The results of detection of PWNs carried by the *Arhopalus rusticus* are shown in table 2. The PWN – carrying beetles accounted for 7.5% of the total. The average number of PWN carried by beetles is 535.

**Table 2.** *Arhopalus rusticus* carry pine wood nematode

| No. <sup>a</sup> | Sex <sup>b</sup> | Number <sup>c</sup> |
| --- | --- | --- |
| 1 | ♀ | 123 |
| 2 | ♀ | 220 |
| 3 | ♀ | 1259 |
| 4 | ♀ | 73 |
| 5 | ♀ | 145 |
| 6 | ♀ | 343 |
| 7 | ♀ | 680 |
| 8 | ♀ | 141 |
| 9 | ♂ | 1450 |
| 10 | ♂ | 433 |
| 11 | ♂ | 518 |
| 12 | ♂ | 171 |
| 13 | ♂ | 1585 |
| 14 | ♂ | 776 |
| 15 | ♂ | 109 |

<sup>a</sup>:serial number of *Arhopalus rusticus*. <sup>b</sup>:The sex of *Arhopalus rusticus*. <sup>c</sup>:The number of pine wood nematodes carried by *Arhopalus rustic*

The population of *Arhopalus rusticus* is large and can carry PWN. If it can transmit PWN, will be a serious hazard to forestry. In order to verify whether *Arhopalus rusticus* transmission PWN, we did the following experiments.

### 3.2 The emergence of *Arhopalus rusticus*

The emergence characteristics of *Arhopalus rusticus* is shown in Fig 2. Three days as a counting unit. The emergence period of *Arhopalus rusticus* is from May 10th to July 11th. The peak period of emergence of *Arhopalus rusticu* was late May and early June, and there was a small emergence peak in late June. In this study, 288 *Arhopalus rusticus* were collected, including 142 female and 146 male beetles. There is no significant difference in the emergence characteristics of the male and female beetles (*P*=0.053).

**Figure 2.**
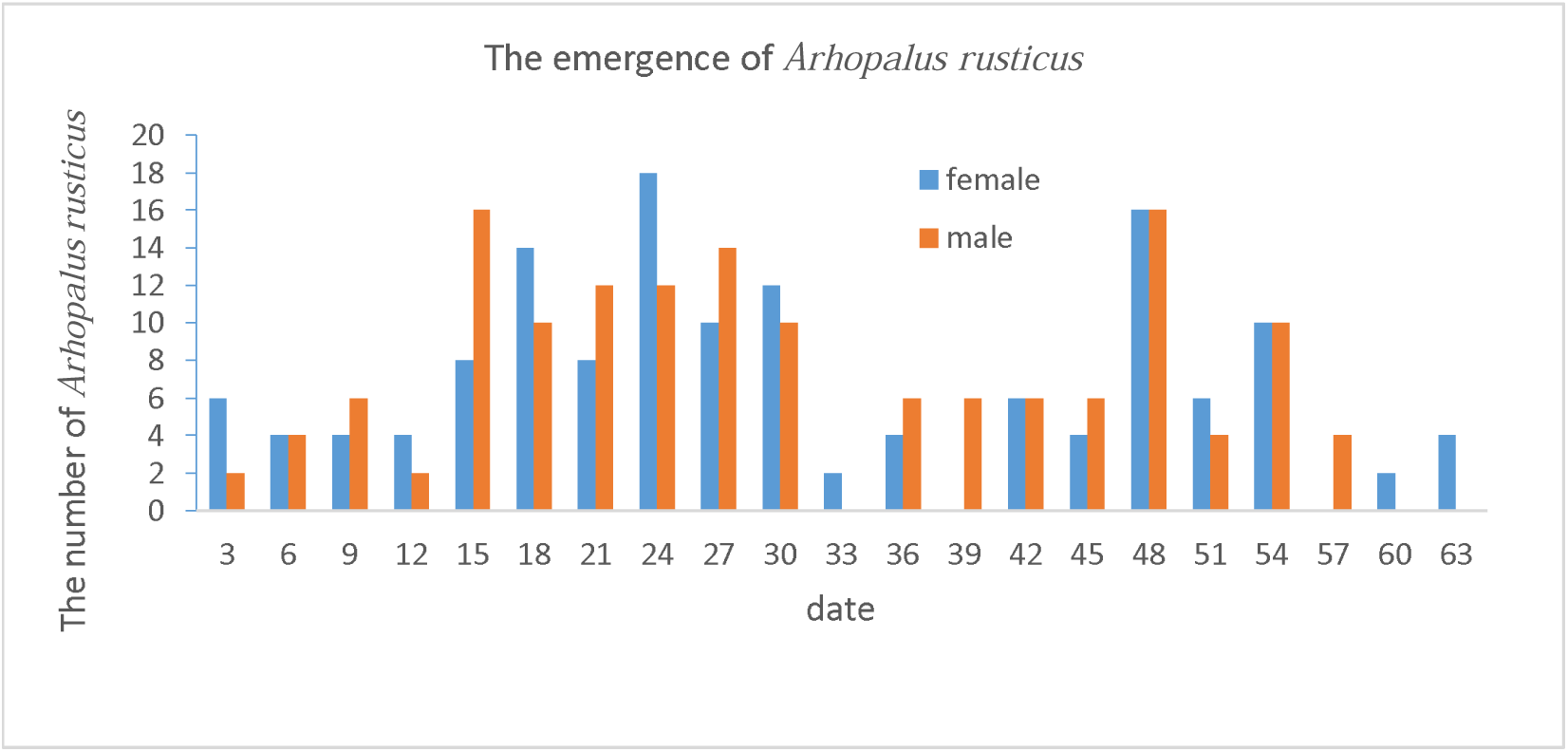
The emergence of Arhopalus rusticus.

### 3.3 The feeding and spawning of the *Arhopalus rusticus*

After emergence, *Arhopalus rusticus* feeding fresh Pine needles to supplement nutrition. There were a large number of obvious bite marks, such as figure 3-D. After the feeding of *Arhopalus rusticus*, the Pine needles gradually wither away from the bite marks and have a distinct boundary as shown in figure 3-A. The fresh bite marks appear emerald green, and can see the obvious turpentine as shown in figure 3-C (arrow), but the pine needles that are withered after feeding are less turpentine as shown in figure 3-B. *Arhopalus rusticus* only caused bite marks on pine needles but did not eat pine needles tissue. It is judged that *Arhopalus rusticus* only absorbs turpentine. This may be related to the structure of the mouthpart (Fig 6), the related characteristics need to be further studied.

**Figure 3.**
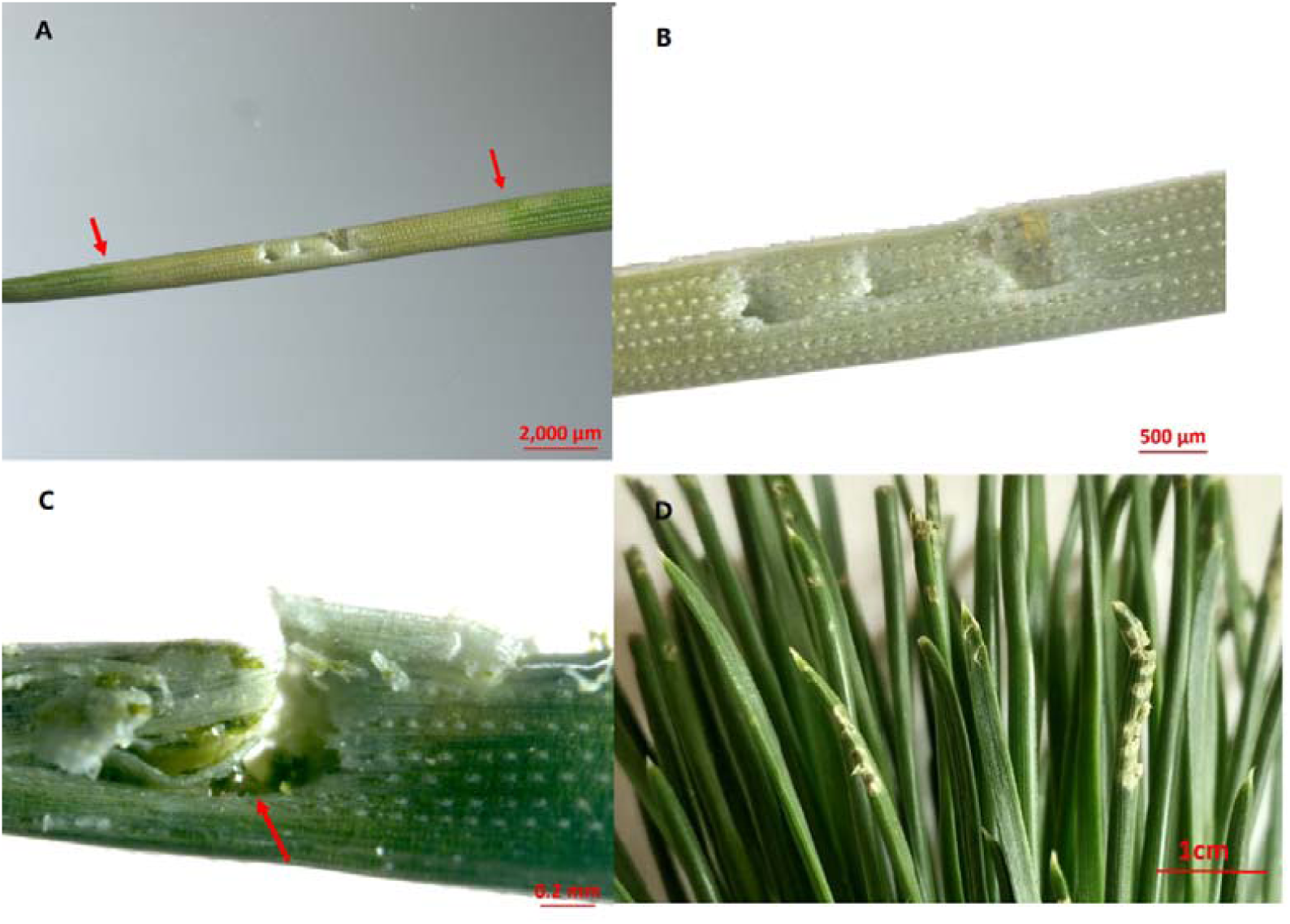
The feeding characteristics of *Arhopalus rusticus*. A, B: The pine needles that withered after the feeding of *Arhopalus rusticus*. C, D: fresh bite marks.

**Figure 4.**
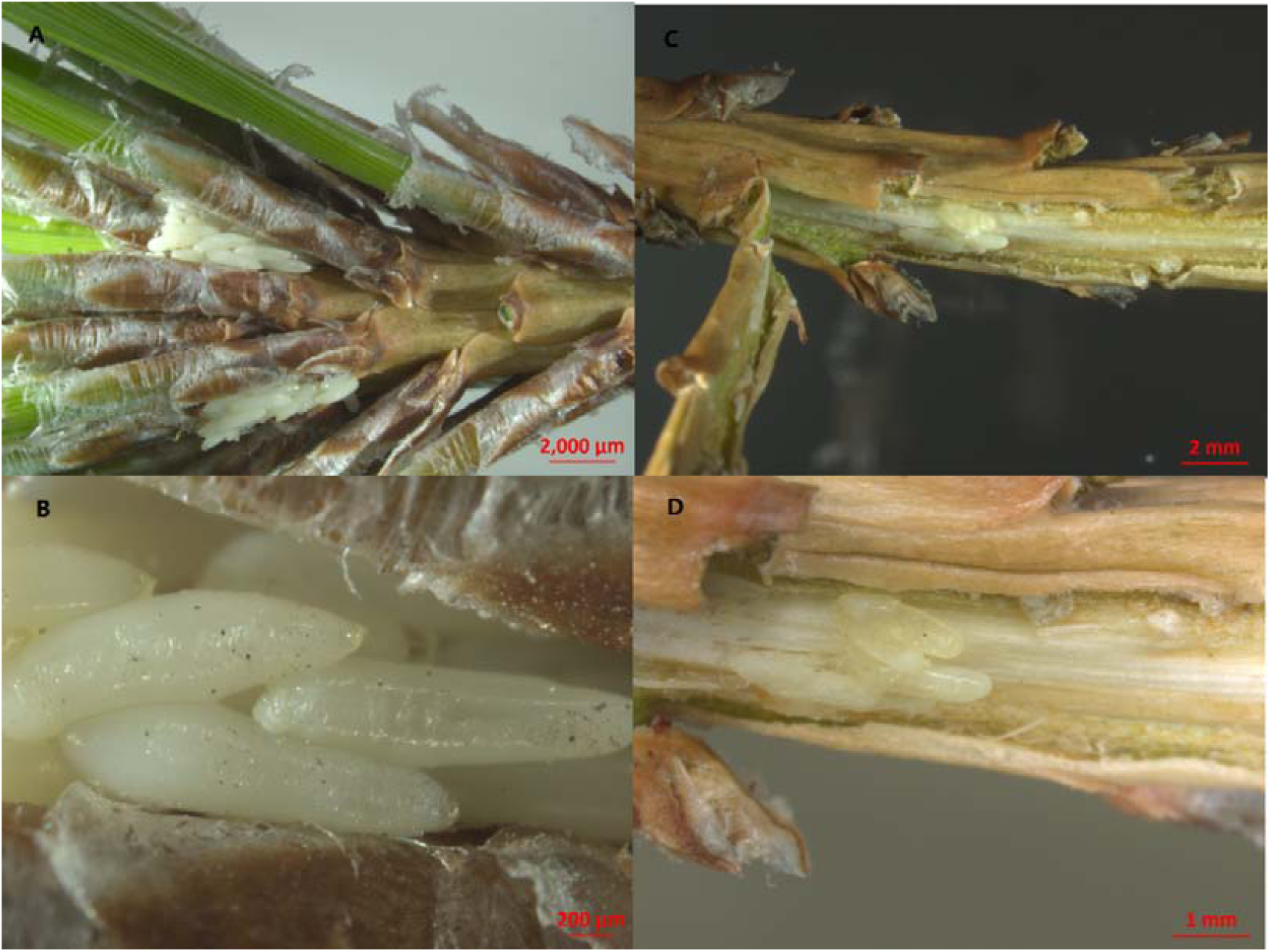
The characteristics of the spawning of *Arhopalus rusticus*.

**Figure 5.**
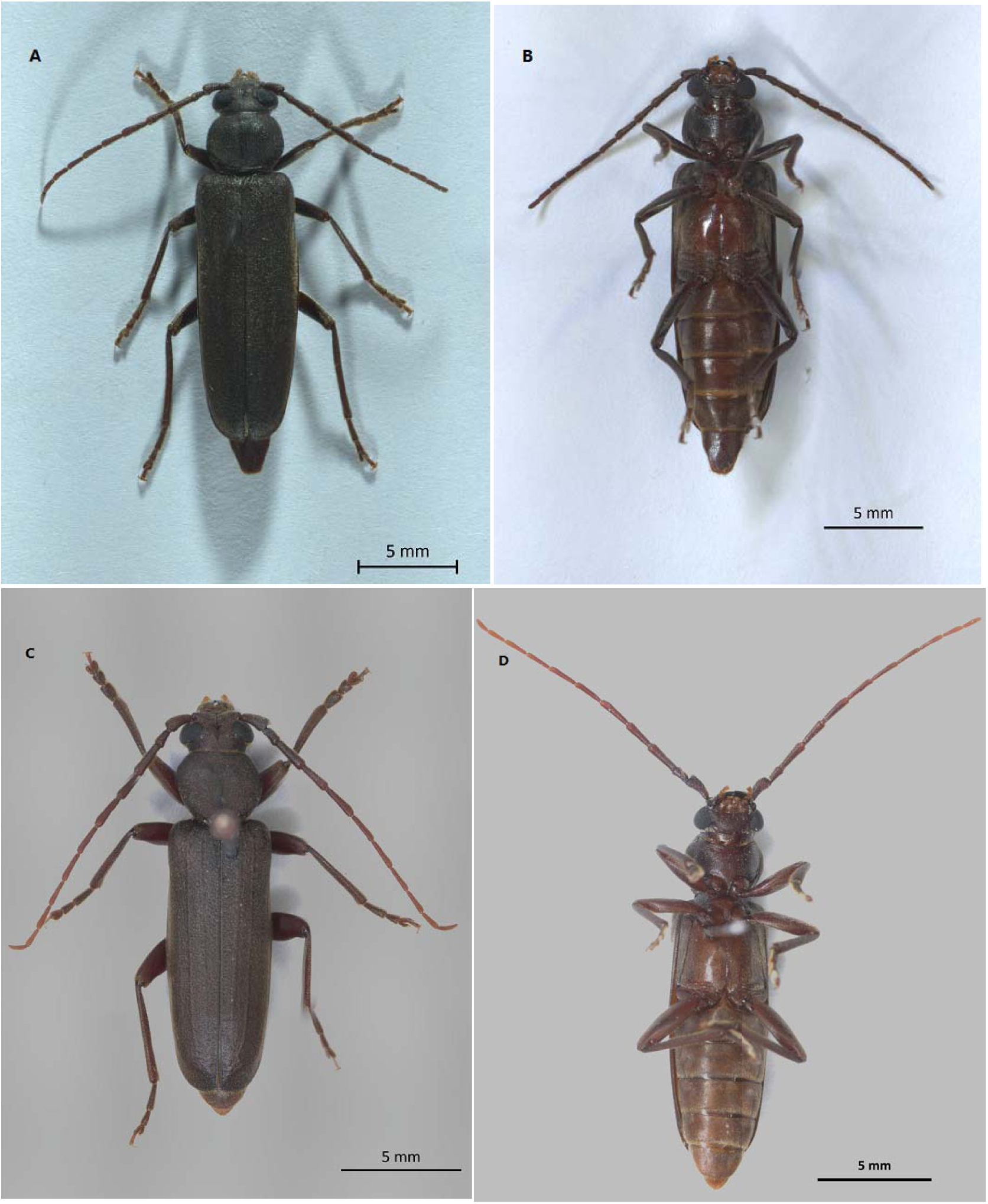
Arhopalus rusticus. A: The back of female *Arhopalus rusticus*. B:The abdomen of female *Arhopalus rusticus*. C: The back of male *Arhopalus rusticus*. D: The abdomen of male *Arhopalus rusticus*.

**Figure 6.**
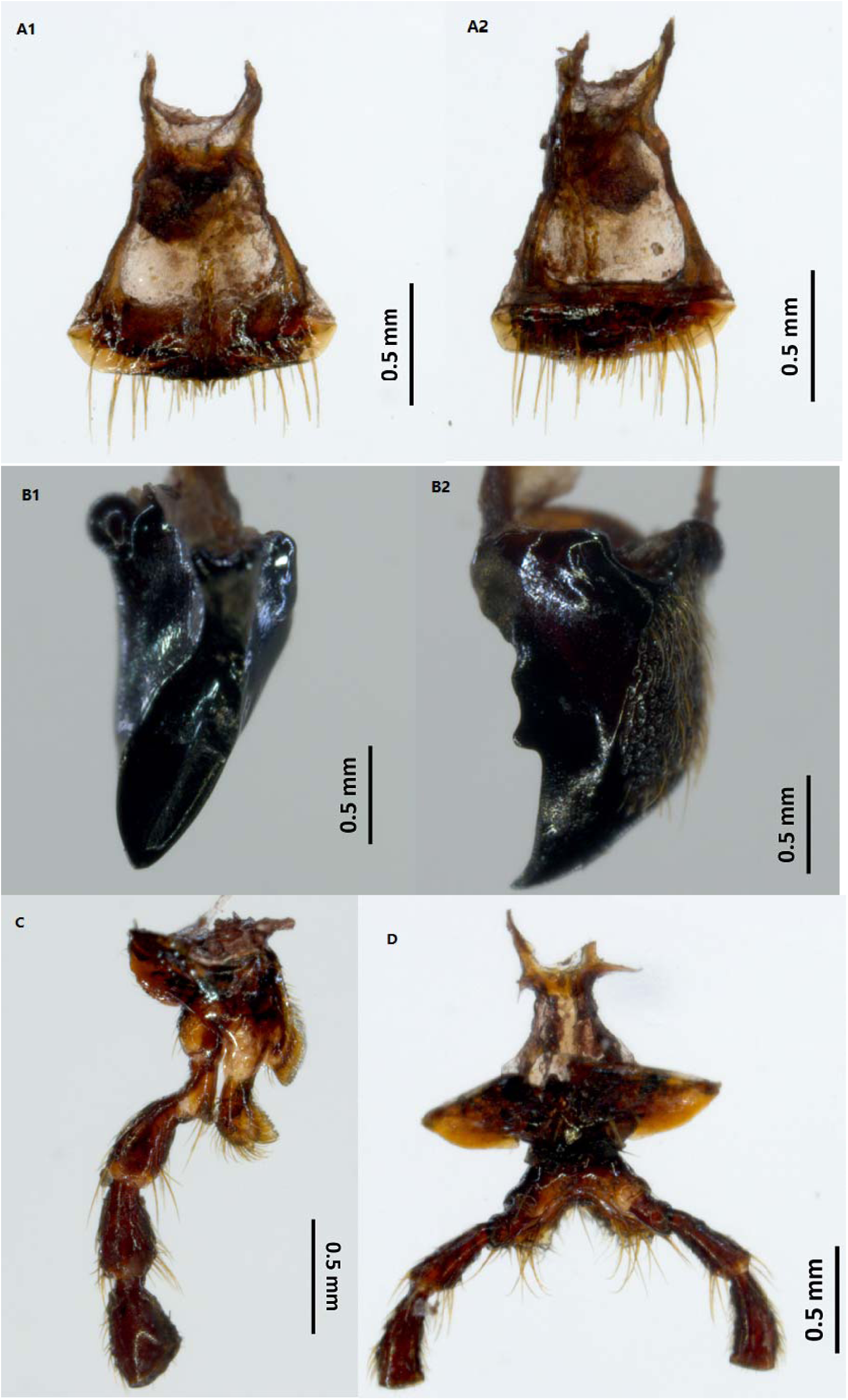
Mouthpart of Arhopalus rusticus. A1, A2: labrum. B1, B2: maxilla. C: mandible. D: Labium

*Arhopalus rusticus* can mate on the day of emergence, and lay eggs the next day after mating. In general, the beetle lays eggs in the roots of pine needles such as Fig 4-A, 4-B, and under the bark as shown in Fig 4-C, 4-D in clusters. Not all female beetles can spawn. In this study, 48 female beetles had an average of 90 eggs. The size of the egg is 2mm×0.4mm.

### 3.4 *Arhopalus rusticus* carrying and spreading pine wood nematode

The characteristics of *Arhopalus rusticus* carrying and spreading of PWN is shown in table 3. It has been identified that 20 beetles can carry pine wood nematode. It can be seen from the table 3 that have 12 of the 20 beetles can transmit PWN, 7 males and 5 females. All beetles that can transmit PWNs can feeding pine needles. It is presumed that PWN could be transmitted through the bite marks of conifers. Female beetles that can transmit PWN not only feeding pine needles, but also spawning. It is presumed that PWN may be transmitted through the oviposition behavior.

**Table 3.** *Arhopalus rusticus* spread pine wood nematode

| No./sex <sup>a</sup> | Number of PWNs<br>Remaining <sup>b</sup> | Feeding <sup>c</sup> | Longevity (d) <sup>d</sup> | Number of PWNs<br>transmitted | Oviposition<br>Number <sup>e</sup> |
| --- | --- | --- | --- | --- | --- |
| 1/♂ | 400 | √ | 8 | 46 | — |
| 2/♂ | 140 | √ | 11 | 0 | — |
| 3/♂ | 1100 | √ | 7 | 140 | — |
| 4/♂ | 342 | √ | 11 | 0 | — |
| 5/♂ | 495 | √ | 10 | 56 | — |
| 6/♂ | 832 | √ | 7 | 68 | — |
| 7/♂ | 400 | √ | 3 | 0 | — |
| 8/♂ | 725 | √ | 12 | 0 | — |
| 9/♂ | 270 | √ | 9 | 112 | — |
| 10/♂ | 721 | √ | 7 | 75 | — |
| 11/♂ | 663 | √ | 5 | 90 | — |
| 12/♂ | 563 | √ | 8 | 55 | — |
| 13/♀ | 1120 | √ | 8 | 125 | 35 |
| 14/♀ | 360 | √ | 3 | 0 | 0 |
| 15/♀ | 215 | √ | 5 | 0 | 26 |
| 16/♀ | 740 | √ | 4 | 40 | 348 |
| 17/♀ | 613 | √ | 8 | 59 | 132 |
| 18/♀ | 287 | √ | 6 | 52 | 92 |
| 19/♀ | 477 | √ | 11 | 0 | 0 |
| 20/♀ | 382 | √ | 10 | 118 | 53 |

<sup>a</sup>: serial number of *Arhopalus rusticus* and sex. <sup>b</sup>: The number of PWN in the body of *Arhopalus rusticus* after the beetle death. <sup>c</sup>: Whether the beetle is taking food. <sup>d</sup>: The Longevity of *Arhopalus rusticus*. <sup>e</sup>: The number of spawn of the *Arhopalus rusticus*.

The average number of PWNs carried by *Arhopalus rusticus* is 594.10. There was no significant difference in the number of PWN carried by female and male beetles(P=0.81). The average number of PWNs transmitted by *Arhopalus rusticus* is 79.69. There was no significant difference in the number of PWN transmitted by female and male beetles(P=0.94). The average longevity of the beetle is 7.65 d.

### 3.5 Experiment of transmit pine wood nematodes through pine needles

This experiment was designed to further verify whether PWN transmit through pine needles. The results of the experiment are shown in table 4. It can be seen from the results that 7 of the 10 repeats can detect PWN in vial bottles, and PWNs were extracted from 9 of the pine needles. It is suggest that PWN can transmit through pine needles.

**Table 4.** The results of the experiment of the transmission of PWN through pine needles

| No. | Number of PWNs transmitted | Number of PWNs in pine needles | The number of worms in tampons |
| --- | --- | --- | --- |
| 1 | 37 | 8 | 1000 |
| 2 | 5 | 12 | 1000 |
| 3 | 15 | 7 | 1000 |
| 4 | 17 | 0 | 1000 |
| 5 | 0 | 13 | 1000 |
| 6 | 14 | 5 | 1000 |
| 7 | 4 | 6 | 1000 |
| 8 | 0 | 4 | 1000 |
| 9 | 0 | 2 | 1000 |
| 10 | 6 | 7 | 1000 |

## 4 Discussion

Studies suggest that vector insects cause wounds on the surface of the twigs by feeding, and pine wood nematodes fall off into the wound, then transmit to the inside of the tree[19-23]. However, in this study, it was not found that the beetles caused wounds on the surface of the twigs. But, it was found through experiments that *Arhopalus rusticus* can feeding pine needles and cause wounds on needles.

It was found in this study that pine wood nematodes could transmit through pine needles. And pine wood nematodes can also be detected from the twigs that feeding *Arhopalus rusticus*. Therefore, we can judge that *Arhopalus rusticus* can transmit pine wood nematodes through pine needles. Pine wood nematodes can be transmitted through pine needles, which has not been found in previous studies. The related mechanisms need to be further studied.

Studies have shown that vector insects can transmit pine wood nematodes by oviposition behavior [23-26]. In this study, PWNs were detected from the female beetle spawning twigs. It can be judged that *Arhopalus rusticus* may be transmit PWN by spawning behavior. The characteristics of *Arhopalus rusticus* transmit PWN by oviposition behavior need further study.

Studies indicate that it took a while after beetles emergence for PWN to transmit. Juveniles exit the tracheal system of beetles within 10 d of the emergence of the beetle from the tree in which it developed[27], at 7 d after emergence beetles start transmit PWN [28], at 10 d after emergence[29]. In this study, the longevity of *Arhopalus rusticus* was only 7.65 d. Some beetles carried PWN but did not transmit, this may be related to a shorter longevity that died before it could began to transmit PWN. The regularity of the transmission of PWN by *Arhopalus rusticus* need to be further studied.

## 5 Conclusion

*Arhopalus rusticus* did not feed on pine bark, after the emergence they feed on pine needles and cause wounds on needles. *Arhopalus rusticus* did not eat the pine needles tissue, but instead eat turpentine. The pine wood nematodes carried by the beetles fell to the wound and into the pine needles. Pine wood nematodes enter the pine branches through resin channels and further damage the pine trees.

## Notes

### Competing Interest Statement

The authors have declared no competing interest.

